# Cetaceans of the Black Sea: where did they survive glacial?

**DOI:** 10.1101/2020.03.02.972463

**Authors:** Davit Dekanoidze, Natia Kopaliani, Zurab Gurielidze, Levan Ninua, Nana Devidze, David Tarkhnishvili

## Abstract

Three species of cetaceans, *Phocoena phocoena*, *Delphinus delphis* and *Tursiops truncatus ponticus* are found in the Black Sea. The Black Sea populations of all three species show morpho-ecological peculiarities that leaded to their subspecific status: *P. p. relicta* (PPR), *D. d. ponticus* (DDP), and *T. t. ponticus* (TTP). It is not clear how long-lasting was their isolation from the core conspecific populations that ensured the development of adaptive features of PPR, DDP, and TTP. The analysis of mitochondrial haplotypes of PPR suggests that the split time of the at least maternal lineage of the Black Sea population of harbour porpoise lasted for over 100 ky (i.e. they should survive at least the latest glacial maximum within the Black Sea). However, the analysis of multiple microsatellite genotypes leaded some authors to suggest that the isolation is much less long, since middle Holocene. We re-analysed published mitochondrial sequences of all three Black Sea cetaceans along with several tens of sequences obtained from the stranded cetaceans. Our analyses suggest that Black Sea populations of all three cetacean species have an important input of populations that survived the last (and maybe earlier) glacial maxima within the Black Sea, most likely in its south-eastern fragment, which did not freeze in winter time even during the glacial peaks. This analysis is supported by both molecular clock approach and simple population modelling based on the assumption on the effective population size range. Different from the PPR, whose Black Sea population is currently fully isolated, there is a limited migration between the Black Sea and Atlantic populations of *T. truncatus* and *D. delphis*, through the Mediterranean “bridge” population. However, the migration rates are not sufficient to overweight differential selection between the Black Sea and Mediterranean populations, and the local morpho-ecological specifics is successfully maintained.

## INTRODUCTION

Three cetacean species present in the Black Sea: bottlenose dolphin (*Tursiops truncatus*); common dolphin (*Delphinus delphis*), and harbour porpoise (*Phocoena phocoena*) (Kleinenberg, 1956). The species are relatively distant phylogenetically; ancestors of genetically closer bottlenose and common dolphins are thought to be split in Pliocene, whereas the ancestors of the porpoise and that of two other species – in early or middle Miocene (Slater et al., 2010; McGowen et.al, 2009). All three species form populations both in the Black Sea and in Atlantic. In Mediterranean basin, harbour porpoise is occasionally found in small groups and it does not appear to form a stable population there, different from common and bottlenose dolphins (Frantzis et al., 2001;Viaud-Martinez et al., 2008; Fontaine et al., 2007; Tonay et al., 2012; Tonay & Dede, 2013; Cucknell et al., 2016).

The Black Sea populations of these species have smaller bodies and show some other morphological differences from the conspecific Atlantic and Mediterranean populations (Kleinenberg 1956; Amaha 1994; Viaud-Martinez et al. 2007, 2008; Galatius & Goldin, 2011). Harbour porpoise and Bottlenose dolphin are genetically distinct, considering their maternal lineages (Viaud-Martinez et al. 2007, 2008; Tonay et al. 2012). Consequently, the Black Sea populations of cetaceans are described as distinct subspecies: *P. phocoena relicta* (PPR; Tzalkin, 1938), *D. delphis ponticus* (DDP; Barabash-Nikiforov, 1940), and *T. truncatus ponticus* (TTP; Barabasch-Nikiforov, 1960). All three subspecies are included in the IUCN Red List under different categories of threat. TTP is included under category EN (A2cde) (Birkun, 2012), DDP under category VU (A2cde) (Birkun & Franzis, 2008), and PPR under category EN (A1d+4cde) (Birkun & Franzis, 2008). It is thought that the population of DDP is relatively numerous and stable, counting few tens of thousands of adult individuals (Birkun, 2008). Population of TTP is much smaller (Birkun, 2008, 2012), and the population of PPR heavily declined in recent years, probably due to extensive hunting (Birkun & Franzis, 2008); current effective size of the Black Sea porpoise should not be exceeding 700 individuals (Fontaine et al., 2012), although the census population size is much higher (Kopaliani et al. 2017).

There is no consensus about the split time between the Black Sea and Atlantic/ Mediterranean populations of Cetaceans. The level of divergence of mitochondrial DNA fragments between the Atlantic and Black Sea populations of harbour porpoise suggests their separation at least for 175 kyr (Tolley & Rosel, 2006; Viaud-Martínez et al., 2007). However, Fontaine et al. (2010, 2012, 2014) disputed this dating, using the Bayesian model of divergence, based on ten microsatellite loci analysis, as an argument.

The inference of Fontain et al. (2012; 2014) is however based on an assumption about strong selection underlining the evolution of mitochondrial genome, and the authors acknowledge this complication in the interpretation of the time of divergence between the Black Sea and the Atlantic population of harbor porpoises. Besides, the Model for Interdisciplinary Research on Climate (MIROC) (Braconnot *et al*., 2007) that was shown to be more realistic than the other palaeoclimatic models (Tarkhnishvili et al., 2012) suggests positive winter temperature at least at the south-eastern Black Sea Coast, which excludes complete freezing of the Black Sea surface and does not exclude presence of cetacean populations within the basin around the year.

To our knowledge, nobody tried to interpret genetic differentiation between DDP and TTP and the conspecific populations of common and bottlenose dolphins from the Mediterranean and Atlantic. Meanwhile, the comparative analysis of the genetic differentiation between the Black Sea, Mediterranean, and Atlantic populations of all three species may provide us with better understanding of time and factors that triggered the development of the isolated Black Sea populations of all cetacean species.

We re-analysed the published mitochondrial sequences of all three cetacean species, *Phocoena phocoena, Delphinus delphis*, and *Tursiops truncatus*, along with the original sequences of the stranded animals collected on Georgia’s Black Sea coast, and inferred the most likely isolation time (and isolation degree) between the Black Sea and the Atlantic populations of these species. Our inference favours the earlier hypothesis of Pleistocenic isolation of the Black Sea populations of harbour porpoise and, respectively, the two other cetacean species. Consequently, the findings of Fontaine et al. (2012; 2014), in our opinion, do not provide evidence for mid-Holocene isolation of PPR and need an alternative interpretation.

## 2. MATERIAL AND METHODS

### 2.1. Matrilineal diversity

In 2010-2017 we collected tissue samples of dolphins stranded at the eastern Black Sea Coast, between the rivers Chorokhi and Enguri in Georgia.Those included 36 *Phocoena phocoena*, 16 *Delphinus delphis*, and 6 *Tursiops truncatus*. DNA has been extracted from the samples following Qiagen tissue kit.

The mitochondrial (mt) DNA control region were amplified using polymerase chain reaction with different primers. For PPR L15824 (5’-CCTCACTCCTCCCTAAGACT-3’) and H16265 (5’-GCCCGGTGCGAGAAGAGG-3’) was used (Rosel et.al., 1999); for DDP 5’-ACACCAGTCTTGTAAACC-3’ and 5’-TACCAAATGTATGAAACCTCA G-3’ was used (Rosel et al., 1994); for TTP DLTurs-r (5’-CCTGAAGTAAGAACCAGATGTCTTATAAA-3’) and DLTurs-f (5’-CCATTCCTCCTAAGACTCAAGGAA-3’’were used (Viaud-Martinez et.al., 2008). PCR was conducted under the following conditions: 20 μL total volume, with 2-4 μm template DNA, 1x pViromega buffer, 1.25 μM MgCl2, 0.1 μM of each primer, 0.1 μM of each dNTP’s and 0.2 U *Taq* DNA polymerase. Thermo cycler profile harbour porpoise: initial denaturation at 95°C for 3 min, followed by 30 cycles of 40 s at 94°C, 60 s at 55°C and 10 s at 72°C, and a final extension at 72°C for 10 min. The thermo cycler profile was the same for bottlenose and common dolphins: initial denaturation at 95°C for 3 min, followed by 40 cycles of 45 s at 94°C, 1 min at 55°C and 1.30 min at 72°C, and a final extension at 72°C for 8 min. An aliquot of 3–5 μl from each PCR was run on a 1% agarose gel to visualize the DNA fragments by SYBRSafe staining. The amplicons were sequenced on the automatic sequencer ABI 3130. Single stranded sequencing was performed with the same primers as used for PCR, and the Big-Dye Terminator v3.1. PCR fragments were sequenced in both directions to assure sequence accuracy. The alignment of the sequences was performed with BioEdit v7.0 (Hall 1999).

The obtained fragments of control region gene (308 base pairs for PPR, 320 base pairs for DDP and 363 base pairs for TTP) were aligned using BioEdit v7.0 (Hall 1999) along with 247 homological sequences of *P. phocoena, D. delphis*, and *T. truncatus*, downloaded from GenBank. The reference numbers of both our and downloaded sequences and relevant publications are shown in Table S1. In total (both our data and downloaded sequences), 178 sequences of *P. phocoena* were studied (54 from the Black Sea and 122 from Atlantic), 73 sequences of *D. delphis* (19 from the Black Sea, 20 from Atlantic, 43 from Mediterranean an 5 from Indian Ocean), 45 sequences of *T. truncatus* (12 from the Black Sea, 10 from Atlantic, and 23 from Mediterranean).

### 2.2. Inferring matrilineal phylogeny of PPR, DDP, and TTP

Alignment of the sequences was performed in BioEdit 7.0 (Hall, 1999). Sequence divergences were estimated using MEGA 5.1 (Kumar, Tamura & Nei, 2004). The 58 high quality sequence fragments were used in the final analyses (36-PPR; 16-DDP; 6-TTP), conducted separately for each species and combining the samples from the Black Sea, Mediterranean, Atlantic and Indian Oceans. Only unique haplotypes were used in this analysis. We counted the number of variable nucleotide positions, tested the molecular clock hypothesis (Hasegawa, Kishino & Yano, 1985), and found the best model of nucleotide substitution using the Bayesian information criterion (BIC) (Tamura *et al*., 2011). A Bayesian phylogenetic analysis was performed using the software BEAST v1.5.1 (Drummond & Rambaut, 2012) to build phylogenetic trees and to estimate the timings of divergence between the geographic populations. Divergence times (average and 0.95 confidence intervals) were estimated by including the estimated time of split from the closest outgroup and the ingroup inferred at www.timetree.org using methodology of Hedges and Kumar (2009) and Hedges et al. (2015) and based on the five independent estimates of split time for *P. phocoena* and *Phocena dalli* and eleven independent estimates of split time for *Tursiops truncatus* and *T. aduncus*. The Bayesian analysis was initiated from random starting trees, assuming the uncorrelated log-normal relaxed clock model and a coalescent model with constant population size. Posterior distributions of parameters were approximated using Markov chain Monte-Carlo (MCMC) with chain settings that provided sufficient sample size for each parameter (i.e. effective sample size (ESS) > 100, specifically length of chain 100,000,000 for each species.

### 2.3. Inferring Fst and migration rates

For inferring pair-wise matrilineal genetic differentiation, based on the analysed of control region sequence fragments of *P. phocoena* from the Black Sea and Atlantic Ocean and for *D. delphis* and *T. truncatus* from the Black Sea, Mediterranean Sea and the Atlantic, we applied AMOVA algorithm (Excoffier et al. 1992; Kayser et al., 2001) using ARLEQUIN v3.5 (Excoffier & Lischer 2010). Based on the obtained Fst values for haploid loci, we calculated the indexes of migration rates using the equation Fst= 1/(1+2Nm) (Wright, 1951). Although this equation is commonly criticized (Witlock & McCauley, 1999), it is suggested that it shows sufficiently well relative measures of gene flow (Neigel, 2002). We used the inferred values as an index of relative gene flow between Black Sea, Mediterranean, and Atlantic populations of the three studied dolphin species.

### 2.4. Infering isolation time from Fst estimate in harbour porpoise

PPR is completely isolated from the Atlantic population of harbour porpoise (Tonay et al., 2012; Fontaine et al.,2012; Fontaine et al.,2014). If two populations are isolated for *t* generations, the genetic differentiation increases with time following the equation Fst = 1 – (1 – 1/2Ne)^*t*^, where Ne is effective population size (Weir, 1997). Considering real generation time in harbor porpoise varying between median value 10 (Birkun & Franzis, 2008) and 11.9 years (Taylor et al., 2007), we developed a diagram connecting Ne and isolation time, given really observed Fst value and discussed the possible time of matrilineal isolation in this context. We used the estimates for obtaining relative values of population size and split time in maternal lineages of two other species.

## 3. RESULTS

### 3.1. Haplotype Tree and isolation time of three subspecies

All haplotypes of PPR belong to a single fully monophyletic clade distinct from the haplotypes of the Atlantic, separated from the rest of the species ca. 309 Kya, and with P < 0.05 over 130 Kya while based on the estimated time by www.timetree.org (Fig. 1 a). In contrast, haplotypes of DDP (Fig 1 b) and TTP (Fig 1 c) were comprised of several monophyletic lineages, separated from their sister lineages of the Atlantic in different times. DDP descend from 4 maternal lineages separated from the Mediterranean and Atlantic populations 40 – 191 Kya, and at least one lineage found exclusively in the Black Sea has been separated in average 144 Kya (with P < 0.05 over ca. 50 Kya) (Fig. 1b, Table 1). TTP also belongs to 4 maternal lineages independently separated from the Mediterranean and Atlantic populations. Two lineages were Black Sea individuals dominate, have been split from the other populations 108-138 Kya (with P < 0.05 over 60 Kya) (Fig. 1c). In conclusion, all three Black Sea populations of cetaceans, fully or partly, descend from the maternal lineages isolated within the Black Sea before the last Glacial Maximum, according to the lower confidence limit of the timetree estimate; however, some matrilineal migration of DDP and TTP within and from the Black Sea occurred in Holocene.

**Fig. 1.**
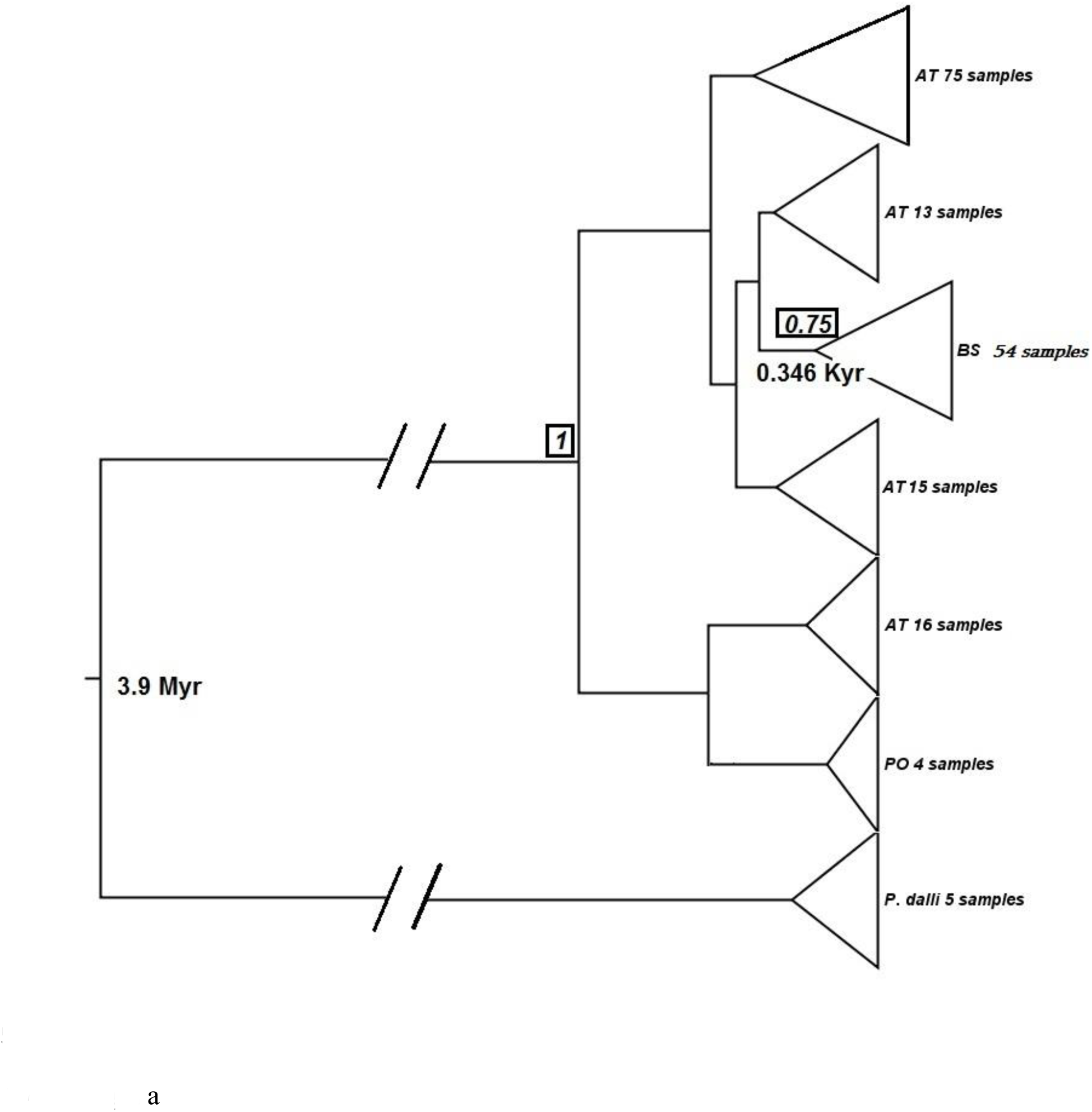
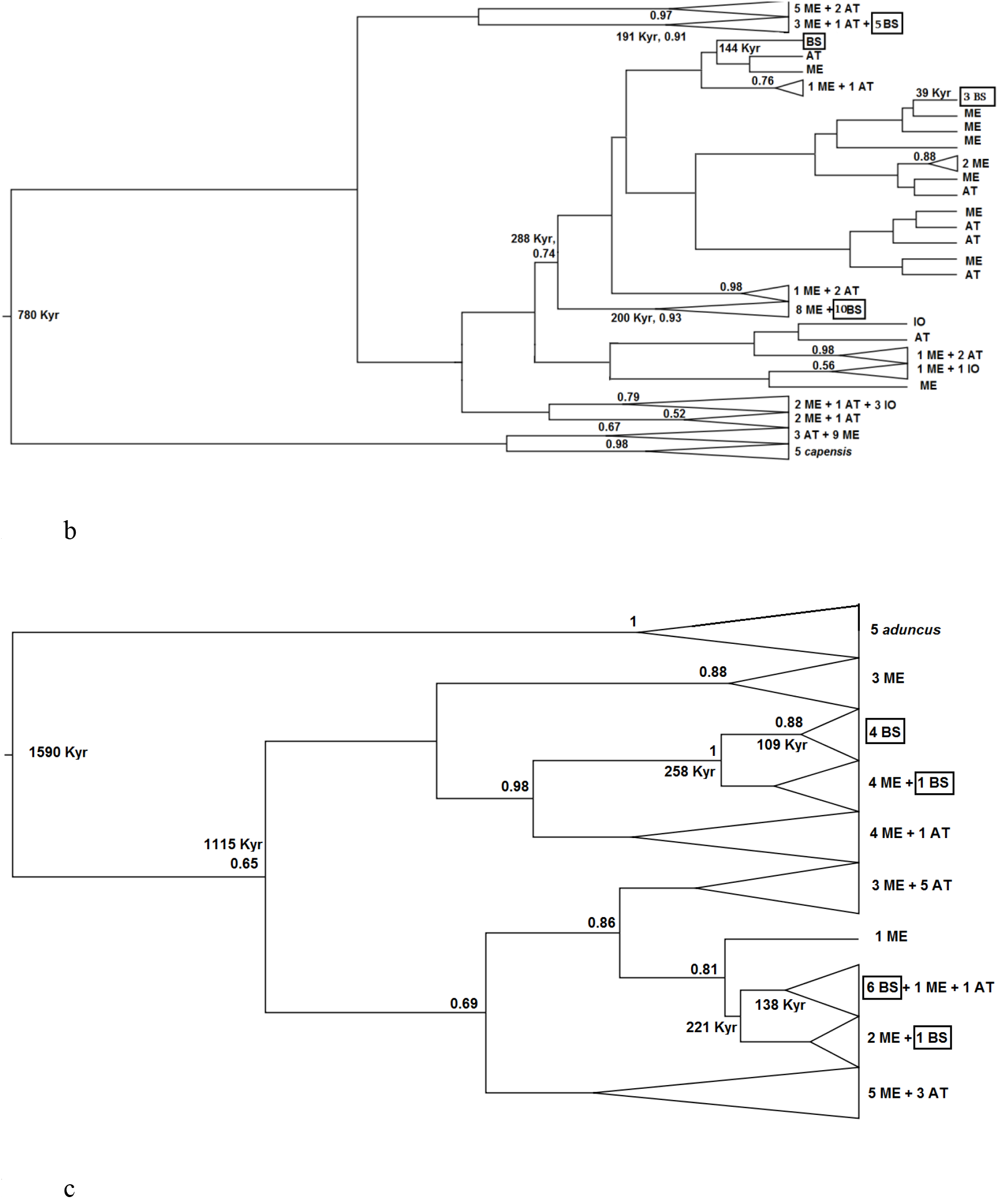
Position of the Black Sea cetaceans within the phylogenetic (Bayesian inference) trees of (a) *Phocoena phocoena*, (b) *Delphinus delphis*, and (c) *Tursiops truncatus*. The list of the reference sequences see in Table S1. Only unique sequences used in this analysis. The numbers with two decimal digits show posterior probabilities for individual clades (only those PPs are shown that exceed 0.5). Kya – thousands of years, the mean age of the clades estimated using calibration of McGowen (2010).

**Table 1.** Average and 95% confidence interval for ages of the cetacean clades found within the Black Sea.

| Taxon | Number of lineages | average split time (Kya), 95% confidence intervals in parenthesis |  |  |  |
| --- | --- | --- | --- | --- | --- |
|  |  | Clade 1 | Clade 2 | Clade 3 | Clade 4 |
| *P. phocoena relicta* | 1 | 309 (136-508) |  |  |  |
| *D. delphis ponticus* | 4 | 191 (66-328) | 144 (50-247) | 39 (14-68) | 171 (59-294) |
| *T. truncatus ponticus* | 4 | 109 (63-156) | 24 (17-41) | 138 (81-198) | 55 (32-79) |

### 3.2. Genetic differentiation among the geographic populations

AMOVA algorithm revealed significant genetic differences between the Black Sea and Atlantic populations for all three species. The differences were higher in common porpoise (Fst=0.46022; P=0.0002, *Nm*=0.586) than in either common dolphin (Fst=0.12557; P=0.0090, *Nm*=3.482) or bottlenose dolphin (Fst=0.12324; P=0.0451, *Nm*=1.779). The differences between the Black Sea and Mediterranean Sea populations of common and bottlenose dolphins were much lower Fst=0.07772, P=0.0360 and Fst=0.04859 (insignificant) than between Black Sea and Atlantic populations of both species. The differences between Mediterranean and Atlantic populations of the same species were Fst=-0.0213 and Fst= 0.03085, both insignificant. Gene flow (*Nm*) between the Black Sea and Atlantic populations of common and bottlenose dolphins is approximately 3.5 individuals per generation, and gene flow between the Black Sea and Mediterranean populations of both species exceeds 15.

### 3.3. Estimation of effective population size vs matrilineal isolation time

The estimated differentiation level in PPR (Fst = 0.46022), given the number of generations ~ 150 (as suggested by Fontaine et al., 2012), considers effective population size varying within 102-122, under condition that generation time is 10-11.9 years. Census population size in mammals usually exceeds effective population size 2-4 times, with few exceptions (Palstra & Fraser, 2012). Hence, the Fst could reach its observed value (0.46) if real population size did not exceed one thousand. This inference is strongly underestimated. The average number of porpoises killed per year in the Black Sea in late 1970-s was nearly 30,000, and although it is considered that in recent years population size of PPR dropped dramatically, the number of wintering porpoises assessed in 2014 just within Georgian territorial waters exceeded 10,000 (Kopaliani et al., unpublished data). Even the minimum (3,464) estimate of Fontaine et al. (2012) for PPR is too high for the observed Fst, if its value is the result of isolation and gene drift. Fontaine et al. (2012) hypothesize selection against mitochondrial genome in this species to explain the observed pattern. With more realistic estimates of effective population size by the same authors (7,518), the number of time required for achieving the observed genetic differentiation would vary between 60,000-170,000 years (Fig. 2).

**Fig. 2.**
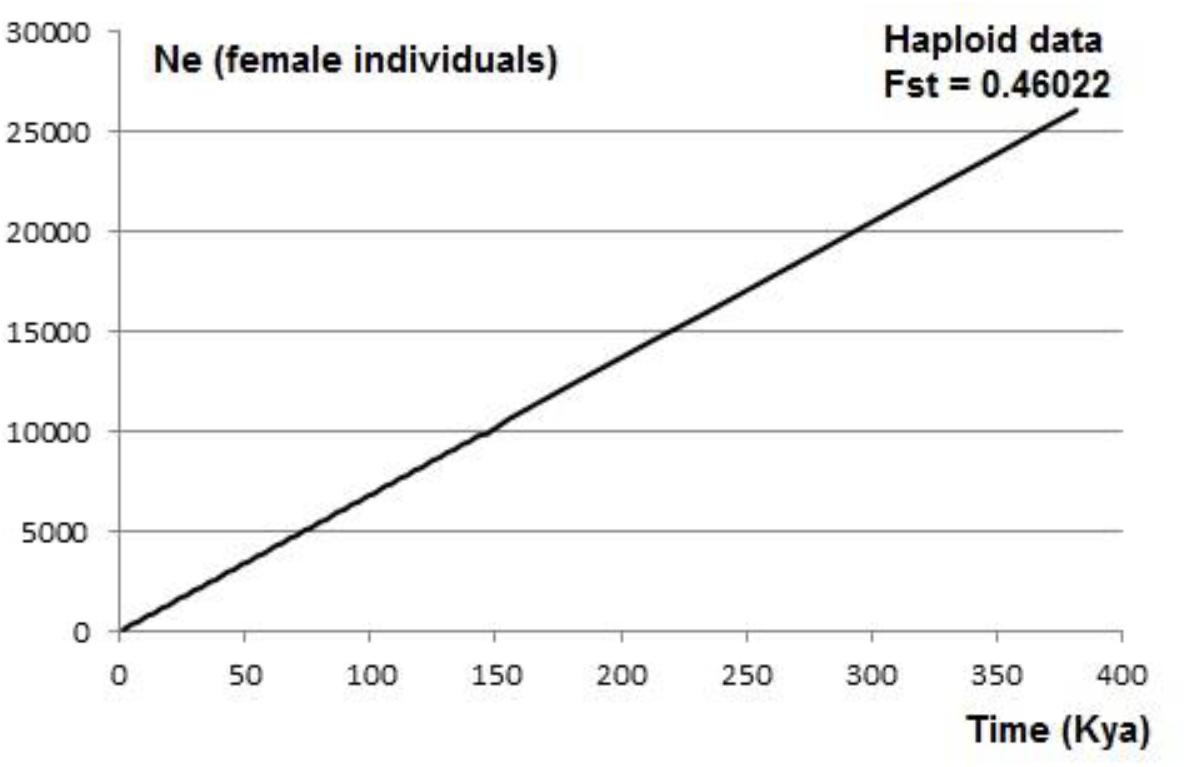
The diagram relating, under the estimated matrilineal differentiation between the Atlantic and Black Sea populations of *P. phocoena* (haploid data Fst = 0.46022), the inferred isolation time (Kya; abscisses) with the effective female population size (ordinates). Generation time considered 11.9 years (Taylor et al., 2007).

## DISCUSSION

All genetic analyses provided both in this paper and earlier papers of (Tolley & Rosel, 2006; Viaud-Martínez et al., 2007) suggest that at least some of the mitochondrial lineages of all three species of cetaceans of the Black Sea descend from the animals that survived the last glacial maximum locally. This fact suggests that, at least in some parts, Black Sea had conditions sufficient for maintaining viable populations of sea mammals. It also explains morphological differences of the Black Sea subspecies of *Tursiops truncatus, Delphinus delphis, and Phocoena phocoena* (Kleinenberg 1956; Amaha 1994; Viaud-Martinez et al. 2007, 2008; Galatius & Goldin, 2011).

The average depth of Bosporus, connecting Black and Mediterranean seas, is about 65 m, and the Black Sea used to be isolated when the sea level was 50-60 m deeper than present. The latest reconnection of the seas occurred ca. 9.5 Kya (Badertscher et al., 2011; Ivanova et al., 2015). Before that, there were relatively long periods of evident water intrusions from the Mediterranean into the Black Sea approx. 60-70, 150-180, 240-270, and 340-360 Kya with shorter intrusion periods in between, and a long period of full isolation ca. 10-50 Kya (Badertscher et al., 2011; Ivanova et al., 2015). The inference reported here suggests that the ancestral lineages of all three Black Sea cetaceans derived during one of the reconnection periods between the Black and Mediterranean Seas and were captured within the Black Sea during consequential glaciation cycles. There is no obvious reason to consider that the cetaceans could not survive long periods in an isolated basin, since multiple fish lineages of Ponto-Caspian fauna, including most likely main prey of PPR, European anchovy (*Engraulis encrasicolus*), survived here for millions of years (Magoulas et al., 1996; Keskin & Atar, 2012). According to recent studies the surface of isolated Black Sea (lake Black) was not covered with permanent ice sheet (Grosswald, 1998; Svendsen et al., 2004). The most plausible model of climate distribution during the Last Glacial Maximum, MIROC (Watanabe et al., 2010; Tarkhnishvili et al., 2012) also suggests positive winter temperature at the eastern Black Sea coast and, hence, absence of full ice cover throughout the Black Sea even in coldest months.

The south-eastern Black Sea coast was an important forest refugium of West Asia and East Europe. This is proved by paleovegetation data (van Andel & Tsedakis, 1996), paleoclimate modeling (Tarkhnishvili et al., 2012; Gavashelishvili & Tarkhnishvili, 2016) and multiple phylogeographic studies (Tarkhnishvili, 2014). Recently, it was shown that this specific area hosted ancestral population of anadromous trout from the entire Black Sea and Caspian drainages (Ninua et al., 2018). That means that not only terrestrial habitats of this area, but also adjacent coastal area of the Black Sea was a critically important aquatic refugium. Given small size and, most likely, low salinity of this refugial sea fragment, one can explain morpho-ecological specifics of the Black Sea cetaceans.

Simultaneously, findings of Fontaine et al. (2010; 2014);2016), suggested much later expansion of the ancestral population of PPR into Black Sea,. One out of three ecotypes of harbor porpoises, Black Sea ecotype split from the other two, European continental shelf waters and upwelling zones of Iberia and Mauritania ecotypes relatively recently (14 kyrbp). The time of divergence between those three ecotypes is the last glacial maximum (23-19kyrbp) and harbor porpoises from the upwelling zones and the BS shared a common ancestor prior to splitting from porpoises currently living on the European continental shelf and (Fontaine et al. 2014). In fact, these findings are based on multiple recombinant microsatellite loci and a 5085 base-pair portion of the mitochondrial genome. For many vertebtrates, there is shown a differential dispersal ability of sexes, with males more easily migrating on large distances than females (Greenwood, 1980; Murtskhvaladze et al., 2010). I.e. mitochondrial and microsatellite DNA analyses revealed highly male-biased dispersal In Dall’s poprpoises (Escorza–Trevino & Dizon, 2000). The model of Fontaine et al. (2010, 2014, 2016) suggest that, indeed, in early and middle Holocene it could be migration between the Black Sea and the Atlantic populations of *Phocoena phocoena*, ceased later due to increased salinity of Mediterranean. The discrepancy between the microsatellite and mitochondrial sequence data would suggest, however, that the migration was sex-biased rather than that it was a selection against some mitochondrial haplotypes. Ancestral Atlantic population split is estimated to have occurred 1325-1855 years ago.The migration from the ancestral Atlantic to the Black Sea population took place, but migration rate is very low.

The migration has also started for two other cetacean species, but for *Delphinus* and *Tursiops* it has never ceased and continues until present. Not surprisingly, the migration rates are the highest for *Tursiops truncatus*, the species with the longest individual range about 1076km and less extent for *Delphinus delphis* with individual range about 500km, smaller than in *T. truncatus* but larger than 300km in *P. phocoena* (Wood, 1998; Genov et al., 2012; Read and Westgate, 2007). Simultaneously, for both these species Mediterranean populations represent a natural bridge between the Black Sea and Atlantic populations, which is not true for *Phocoena phocoena*.

Finally one should suggest that a limited past and/or current migration rates between the Atlantic, Mediterranean, and Black Sea populations of cetaceans are insufficient for melting morpho-ecological differences of PPR, DDP and TTP gained during long-lasting adaptive selection of these taxa in specific conditions of limited almost freshwater basin in the south-eastern part of the Black Sea.

## Acknowledgements

The authors thank Ilia State University administration and Kolkheti Protected Areas Development Fund for administrative and financial support; Mari Murtskhvaladze for assistance at the molecular genetic laboratory at Ilia State University and the crew of the Ilia State University research vessel “Saint Ilia” for the help during marine surveys.

## Supplementary materials

**Table S1.** The reference numbers of original and downloaded sequences and relevant publications

| Numbers | Authors | Publication Title |
| --- | --- | --- |
| AB499064 -<br>AB499073 | Taguchi,M., Chivers,S.J.,<br>Rosel,P.E., Matsuishi,T.<br>and Abe,S. | Mitochondrial DNA phylogeography of<br>the harbour porpoise *Phocoena phocoena*<br>in the North Pacific (2010) |
| GQ338845 –<br>GQ338858 | Tolley,K.A., Vikingsson,G.A.<br>and Rosel,P.E. | Mitochondrial DNA sequence variation<br>and phylogeographic patterns in harbour<br>porpoises *Phocoena phocoena* from the<br>North Atlantic (2001) |
| FJ214731 –<br>FJ214801 | Rosel,P.E., France,S.C.,<br>Wang,J.Y. and Kocher,T.D. | Genetic structure of harbour porpoise<br>*Phocoena phocoena* populations in the<br>northwest Atlantic based on mitochondrial<br>and nuclear (1999) |
| EF063110,<br>EF063646 –<br>EF063676 | Viaud-Martinez,K.A.,<br>Martinez Vergara,M.,<br>Gol'din,P.E., Ridoux,V.,<br>Ozturk,A.A., Ozturk,B.,<br>Rosel,P.E., Frantzis,A.,<br>Komnenou,A. and<br>Bohonak,A.J. | Morphological and genetic differentiation<br>of the Black Sea harbor porpoise<br>(*Phocoena phocoena relicta*) (2007) |
| AF461818 –<br>AF461891 | Chivers,S.J., Dizon,A.E.,<br>Gearin,P.J. and<br>Robertson,K.M. | Small-scale population structure of<br>eastern North Pacific harbor, *Phocoena*<br>*phocoena*, indicated by molecular genetic<br>analyses (2002) |
| AF311924 –<br>AF311933 | Tolley,K.A., Vikingsson,G.A.<br>and Rosel,P.E. | Mitochondrial DNA sequence variation<br>and phylogeographic patterns in harbour<br>porpoises *Phocoena phocoena* from the<br>North Atlantic (2001) |
| AF152570 –<br>AF152571 | Rosel,P.E., France,S.C.,<br>Wang,J.Y. and Kocher,T.D. | Genetic structure of harbour porpoise<br>*Phocoena phocoena* populations in the<br>northwest Atlantic based on mitochondrial<br>and nuclear markers (1999) |
| Y13872 –<br>Y13880 | Tiedemann,R., Harder,J.,<br>Gmeiner,C. and Haase,E. | Mitochondrial DNA sequence patterns of<br>Harbour porpoises (*Phocoena phocoena*)<br>from the North and the Baltic Sea (1996) |
| HQ223475 –<br>HQ223479 | Moller,L., Valdez,F.P.,<br>Allen,S., Bilgmann,K.,<br>Corrigan,S. and<br>Beheregaray,L.B. | Fine-scale genetic structure in short-beaked common dolphins *Delphinus*<br>*delphis*) along the East Australian Current<br>(2011) |
| KF281698 | Jung,J.-I., Meheust,E.,<br>Alfonsi,E. and Hassani,S | The use of DNA barcoding to monitor the<br>marine mammal biodiversity along the<br>French Atlantic coast (Unpublished) |
| AF242199 | Canadas,A., Vaquero,C.,<br>Fernandez,E., Sagarminaga,R.,<br>Herranz,M., Marin,A.,<br>Morales,A., Gutierrez,G. and<br>Fernandez-Piqueras,J. | Mitochondrial DNA analysis of common<br>dolphins (*Delphinus delphis*) from<br>Alboran and Black seas (Unpublished) |
| DQ520112 –<br>DQ520124 | Hildebrandt,S., Brown,R.P.,<br>Zamorano Serrano,M.J., Lopez<br>Jurado,L.F. and Afonso<br>Lopez,J.M. | Genetic characterization of *Delphinus*<br>*delphis* from the Canary Islands based on<br>the mitochondrial DNA control region<br>(Unpublished) |
| DDU02639 –<br>DDU02641 | Rosel,P.E., Dizon,A.E. and Heyning,J.E. | Genetic analysis of sympatric morphotypes of common dolphins (genus *Delphinus*)(1994) |
| DQ378113 –<br>DQ378117 | Amaral,A.R., Sequeira,M., Martinez-Cedeira,J. and Coelho,M.M. | New insights on population genetic structure of *Delphinus delphis* from the northeast Atlantic and phylogenetic relationships within the genus inferred from two mitochondrial markers (2007) |
| EU365129 –<br>EU365173 | Natoli,A., Canadas,A., Vaquero,C., Politi,E., Fernandez-Navarro,P. and Hoelzel,A.R. | Conservation genetics of the short-beaked common dolphin (*Delphinus delphis*) in the Mediterranean Sea and in the eastern North Atlantic Ocean (2008) |
| AY963588 –<br>AY963626 | Natoli,A., Birkun,A., Aguilar,A., Lopez,A. and Hoelzel,A.R | Habitat structure and the dispersal of male and female bottlenose dolphins (*Tursiops truncatus*) (2005) |
| Delphinus delphis | David Dekanoidzea; Natia Kopaliania; Zurab Gurielidzea,b; Levan Ninuaa; Nana Devidzea and David Tarkhnishvili | This paper |
| KX364697 -<br>KX364701 |  |  |
| Phocoena phocoena |  |  |
| KX364702 -<br>KX364706 |  |  |
| Tursiops truncatus |  |  |
| KX364707 -<br>KX364711 |  |  |

